# Autoencoder-based cluster ensembles for single-cell RNA-seq data analysis

**DOI:** 10.1101/773903

**Authors:** Thomas A Geddes, Taiyun Kim, Lihao Nan, James G Burchfield, Jean YH Yang, Dacheng Tao, Pengyi Yang

**Affiliations:** Charles Perkins Centre, School of Mathematics and Statistics, Faculty of Science, The University of Sydney, NSW 2006, Australia; UBTECH Sydney Artificial Intelligence Centre and the School of Computer Science, Faculty of Engineering and Information Technologies, The University of Sydney, NSW 2006, Australia; Charles Perkins Centre, School of Life and Environmental Sciences, Faculty of Science, The University of Sydney, NSW 2006, Australia; Computational Systems Biology Group, Children’s Medical Research Institute, Faculty of Medicine and Health, The University of Sydney, NSW 2145, Australia

**Keywords:** Autoencoder, Cluster ensemble, Single cells, scRNA-seq, Single-cell transcriptome, Cell type identification

## Abstract

**Background:** Single-cell RNA-sequencing (scRNA-seq) is a transformative technology, allowing global transcriptomes of individual cells to be profiled with high accuracy. An essential task in scRNA-seq data analysis is the identification of cell types from complex samples or tissues profiled in an experiment. To this end, clustering has become a key computational technique for grouping cells based on their transcriptome profiles, enabling subsequent cell type identification from each cluster of cells. Due to the high feature-dimensionality of the transcriptome (i.e. the large number of measured genes in each cell) and because only a small fraction of genes are cell type-specific and therefore informative for generating cell type-specific clusters, clustering directly on the original feature/gene dimension may lead to uninformative clusters and hinder correct cell type identification.

**Results:** Here, we propose an autoencoder-based cluster ensemble framework in which we first take random subspace projections from the data, then compress each random projection to a low-dimensional space using an autoencoder artificial neural network, and finally apply ensemble clustering across all encoded datasets for generating clusters of cells. We employ four evaluation metrics to benchmark clustering performance and our experiments demonstrate that the proposed autoencoder-based cluster ensemble can lead to substantially improved cell type-specific clusters when applied with both the standard *k*-means clustering algorithm and a state-of-the-art kernel-based clustering algorithm (SIMLR) designed specifically for scRNA-seq data. Compared to directly using these clustering algorithms on the original datasets, the performance improvement in some cases is up to 100%, depending on the evaluation metrics used.

**Conclusions:** Our results suggest that the proposed framework can facilitate more accurate cell type identification as well as other downstream analyses. The code for creating the proposed autoencoder-based cluster ensemble framework is freely available from https://github.com/gedcom/autoencoder_cluster_ensemble

## Background

Transcriptome profiling by single-cell RNA-sequencing (scRNA-seq) is a fast-emerging technology for studying complex tissues and biological systems at the single-cell level [1]. Identification of cell types present in a biological sample or system is a vital part of scRNA-seq data analysis workflow [2]. The key computational technique for unbiased cell type identification from scRNA-seq data is unsupervised clustering [3]. Typically, this is achieved by using a clustering algorithm to partition cells in a scRNA-seq dataset into distinct groups and subsequently annotate each group to a type of cell based on cell type marker genes and/or other biological knowledge of cell type characteristics [4].

Due to the critical role played by cell type identification for downstream analyses, significant amount of effort has been devoted to tailoring standard clustering algorithms or developing new ones for scRNA-seq data clustering and cell type identification [5]. These include standard *k*-means clustering, hierarchical clustering, and variants that are specifically designed for scRNA-seq data (i.e. RaceID/RaceID2 [6], CIDR [7]) as well as more advanced methods that utilise likelihood-based mixture modelling (countClust) [8], density-based spatial clustering [9] and kernel-based single-cell clustering (SIMLR) [10]. Several studies have compared and summarised various clustering algorithms used for scRNA-seq data analysis [11, 12, 13].

One of the key challenges in scRNA-seq data clustering is handling specific characteristics of the data including high feature-dimensionality and high feature-redundancy. This is because that typically a large number of genes are profiled in the experiment but only a small proportion of them are cell type-specific and therefore informative for cell type identification. Hence, clustering directly on the original high-dimensional feature space may result in suboptimal partitioning of the cells due to low signal-to-noise ratio. To reduce the high feature-dimensionality in scRNA-seq data for visualisation and downstream analyses, various dimension reduction techniques, including traditional approaches as well as newly developed ones, have been applied to scRNA-seq data. These include generic methods such as principal component analysis (PCA), independent component analysis (ICA), non-negative matrix factorization [14], and t-distributed stochastic neighbour embedding (tSNE) [15], as well as other methods developed for scRNA-seq data, such as zero inflated factor analysis (ZIFA) [16]. Recently, deep learning techniques such as scvis, a deep generative model [17], and a scNN [18], a neural network model, were developed specifically for scRNA-seq data dimension-reduction. While these new developments are primarily focused on scRNA-seq data visualisation, they represent the first applications of deep learning techniques for scRNA-seq data analysis.

Ensemble learning is an established field in machine learning and has a wide application in bioinformatics [19]. Ensemble clustering via random initialisation is a popular ensemble learning method for clustering [20]. While this approach was found to improve stability of the *k*-means clustering algorithm, it appeared to have a less consistent effect on clustering accuracy [21]. Ensemble clustering via random projection is an alternative ensemble learning method for clustering. This approach was applied to DNA microarray data analysis and resulted in improved clustering accuracy [22]. Weighted ensemble clustering combines multiple clustering outputs based on their respective quality [23]. Recently, cluster ensembles have been generated by combining outputs from different upstream processing and similarity metrics [24] or different clustering algorithms for cell type identification from scRNA-seq data [25, 26]. While these heuristic methods were found to be effective for improving clustering accuracy in cell type identification, they are ad-hoc in nature and may not fully explore characteristics and biological signals in scRNA-seq data when clustering.

To extract biological signal from scRNA-seq data while at the same time addressing the issues of high feature-dimensionality and high feature-redundancy, here we propose an autoencoder-based cluster ensemble framework for the robust clustering of cells for cell type identification from scRNA-seq data. The proposed framework first randomly projects the original scRNA-seq datasets into sub-spaces to create ‘diversity’ [27] and then trains autoencoder networks to compress each such random projection to a low-dimensional feature space. Subsequently, clusterings are generated on all encoded datasets and consolidated into a final ensemble output.

The proposed framework of cluster ensemble via autoencoder-based dimension-reduction and its application to scRNA-seq is a principled approach and the first of its kind. We demonstrate that (1) the autoencoder-reduced ensemble clustering of scRNA-seq data significantly improves clustering accuracy of cell types, whereas simple ensemble clustering without autoencoder-based dimension reduction showed no clear improvement; (2) improvement of clustering accuracy in general increases with the ensemble size; and (3) the proposed framework can improve cell type-specific clustering when applied using either the standard *k*-means clustering algorithm or a state-of-the-art kernel-based clustering algorithm (SIMLR) [10] specifically designed for scRNA-seq data analysis. This demonstrates that the proposed framework can be coupled with different clustering algorithms to facilitate accurate cell type identification and other downstream analyses.

## Results

### Hyperparameter optimisation for autoencoders

We undertook a grid search to optimise three hyperparameters including random projection size, encoded feature space size and autoencoder learning rate during backpropagation; this was performed across four datasets (Table 2) using the ARI, NMI, FM and Jaccard index metrics discussed above. Together, the four metrics across four datasets made a total of sixteen dimensions (i.e.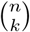) across which to optimise. We used Pareto analysis [29] to select an appropriate combination of parameters from across all four optimisation datasets without giving priority to any single dataset or metric. A Pareto rank of 1 indicates an optimal clustering results on a selection of optimisation datasets using a combination of hyperparameters.

**Table 1.** Comparison of direct application of *k*-means and SIMLR clustering on raw gene expression data with autoencoder-based *k*-means and SIMLR ensemble. Cell type identification accuracy were quantified by the four evaluation metrics.

|  | *k*-means |  |  |  |  |  |  |  |
| --- | --- | --- | --- | --- | --- | --- | --- | --- |
|  | Raw |  |  |  | Autoencoder |  |  |  |
|  | ARI | NMI | FM | Jaccard | ARI | NMI | FM | Jaccard |
| Mouse neurons | 0.22±0.02 | 0.36±0.03 | 0.39±0.02 | 0.22±0.01 | 0.38±0.02 | 0.56±0.01 | 0.53±0.02 | 0.34±0.02 |
| Mouse striatum | 0.36±0.07 | 0.69±0.06 | 0.51±0.06 | 0.31±0.05 | 0.45±0.01 | 0.75±0.01 | 0.58±0.01 | 0.37±0.01 |
| Human archived brain | 0.29±0.01 | 0.49±0.01 | 0.35±0.01 | 0.21±0.01 | 0.37±0.01 | 0.56±0 | 0.43±0.01 | 0.27±0.01 |
| Mouse archived brain | 0.32±0.01 | 0.49±0 | 0.35±0.01 | 0.21±0.01 | 0.43±0.01 | 0.58±0 | 0.46±0.01 | 0.3±0.01 |
|  | SIMLR |  |  |  |  |  |  |  |
|  | Raw |  |  |  | Autoencoder |  |  |  |
|  | ARI | NMI | FM | Jaccard | ARI | NMI | FM | Jaccard |
| Mouse neurons | 0.44±0 | 0.65±0 | 0.58±0 | 0.39±0 | 0.71±0 | 0.7±0 | 0.81±0 | 0.67±0 |
| Mouse striatum | 0.55±0 | 0.81±0 | 0.67±0 | 0.34±0 | 0.8±0.02 | 0.87±0.01 | 0.85±0.02 | 0.74±0.03 |

**Table 2.** Summary of the experimental scRNA-seq datasets used for hyperparameter optimisation and method evaluation.

| Repository | Source | # cell | # class | Ref. | Protocol | Purpose |
| --- | --- | --- | --- | --- | --- | --- |
| GSE60361 | Mouse cortex | 3005 | 7 | [32] | SMARTer | Optimisation |
| GSE45719 | Mouse embryogenesis | 300 | 8 | [33] | SMART-seq2 | Optimisation |
| GSE67835 | Adult and fetal human brain | 466 | 8 | [34] | SMARTer | Optimisation |
| E_MTAB_3929 | Human embryogenesis | 1529 | 5 | [35] | SMART-seq2 | Optimisation |
| GSE84371 | Mouse neurons | 1402 | 8 | [36] | Smart-seq2 | Evaluation |
| GSE82187 | Mouse striatum | 705 | 10 | [37] | SMARTer & Smart-seq2 | Evaluation |
| Broad portal | Human archived brain | 14963 | 19 | [38] | Drop-seq | Evaluation |
| Broad portal | Mouse archived brain | 13313 | 26 | [38] | Drop-seq | Evaluation |

As the Pareto front becomes larger and more ambiguous as more datasets are included, we tested the robustness of each parameter set by obtaining the Pareto front for all possible combinations of 1, 2, 3 or all 4 datasets and counting the number of such combinations for which the given parameter set appears in the Pareto front (Figure 2). We determined that the most robust high-accuracy results were obtained by selecting 2048 genes during random projection; producing an encoded feature space of 16 dimensions; and training the autoencoder using a learning rate of 0.001. All evaluation benchmarks were undertaken using this parameter combination and a hidden layer width of 128.

**Figure 1.**
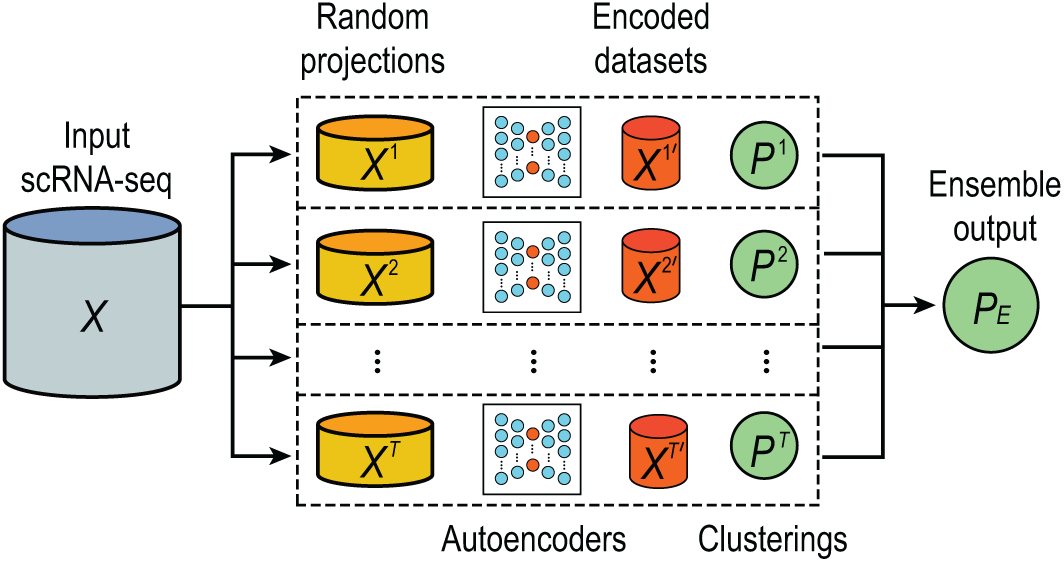
A schematic illustration of the proposed autoencoder-based cluster ensemble framework. The first step is the sampling of multiple random projections from the original input scRNA-seq data set. A separate autoencoder artificial neural network is trained on each of these random projections and used to encode the data to a smaller-dimensional space. Subsequently, clustering of each encoded dataset is conducted using an arbitrary clustering method; the final clustering output is produced by integrating individual clustering results using a fixed-point algorithm [28].

**Figure 2.**
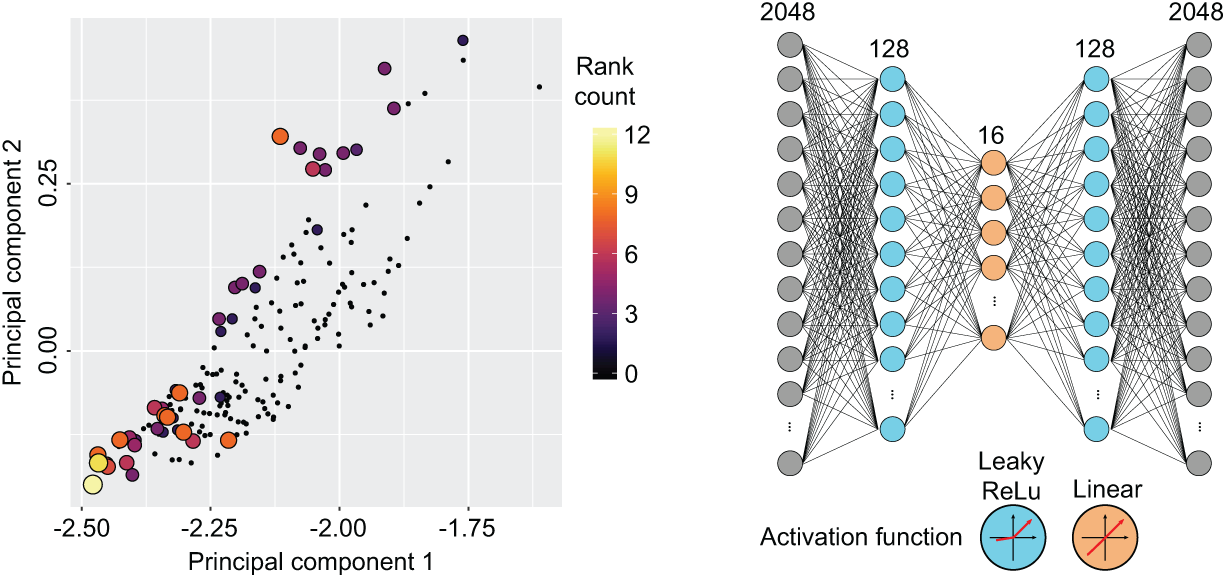
Hyperparameter optimisation for autoencoders using Pareto analysis. Left panel: PCA visualisation of the four evaluation metrics (i.e. ARI, NMI, FM and Jaccard) on each of the four optimisation datasets. Each point corresponds to a single combination of hyperparameter values including random projection size, encoded feature space size, and autoencoder learning rate during backpropagation; each combination/point is colour-coded by the number of times it was assigned Pareto rank 1 (i.e. the combination that gives best clustering performance) across all possible combinations of the four optimisation datasets. Right panel: Autoencoder architecture as determined by the hyperparameter optimisation procedure.

### Ensemble of *k*-means clustering

We first asked if the ensemble of autoencoder-based clustering can improve upon the performance of a single clustering run on a single encoded dataset. To test this, we first used a standard *k*-means clustering algorithm (section) to create base clustering results and tested the performance of different ensemble sizes based on ARI, NMI, FM and Jaccard. Note that we repeatedly run the entire procedure multiple times to account for the variability in the clustering results. We found that in general the overall ensemble clustering performance improves as the number of base clustering runs increases (Figure 3, light blue boxes) according to all four evaluation metrics and in all four datasets used for evaluation. These results demonstrate that the ensemble of autoencoder-based clustering framework indeed can improve cell type identification for the *k*-means clustering algorithm.

**Figure 3.**
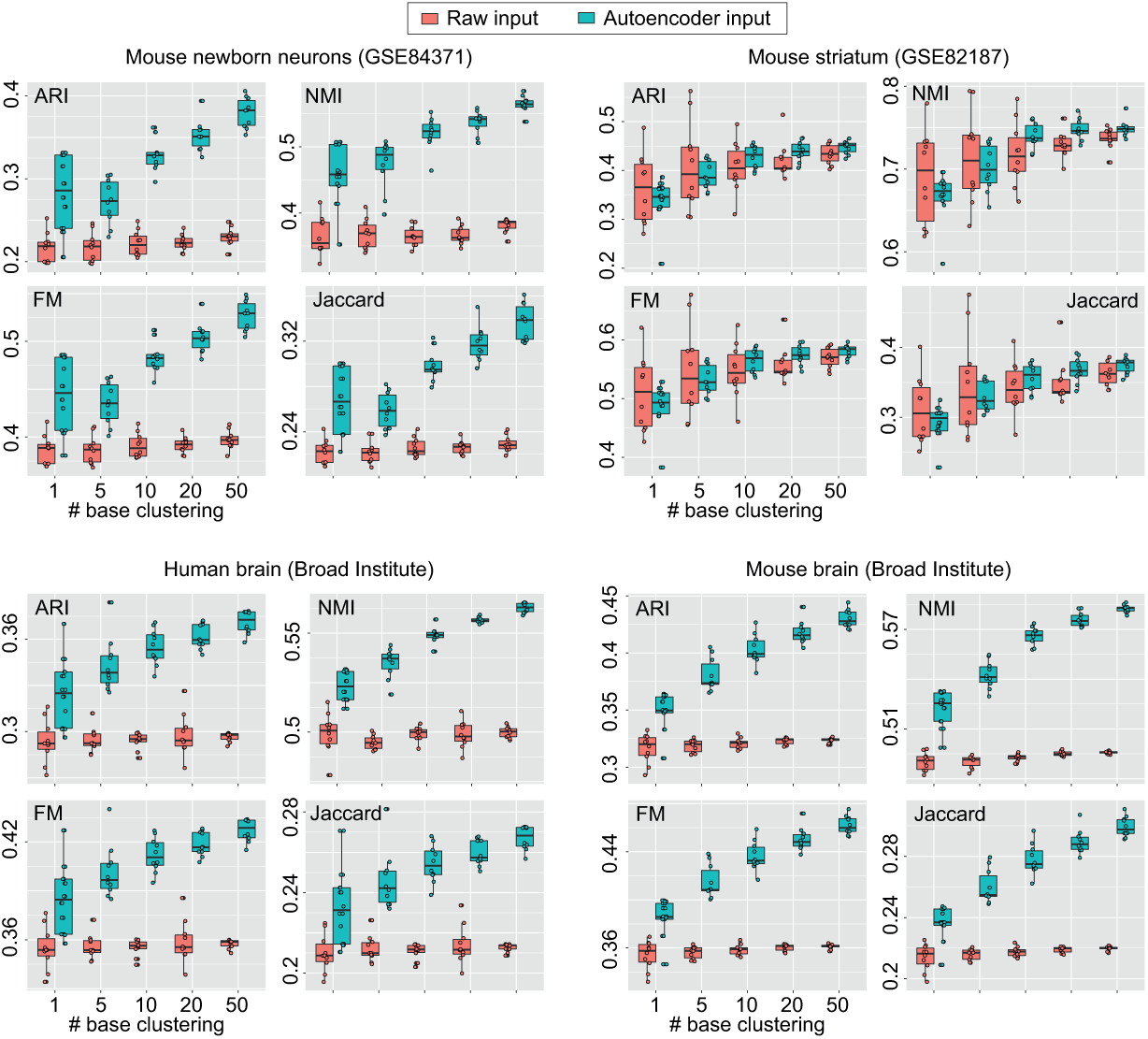
Ensemble of *k*-means clustering results on the four scRNA-seq datasets. Red boxes represent ensemble of *k*-means clustering on the raw input expression matrix without using the autoencoder framework. Light blue boxes represent autoencoder-based *k*-means cluster ensemble.

We wondered if such an improvement of cluster ensemble is independent of the random projection and autoencoder framework. Hence, we compared the performance of *k*-means clustering on the raw input expression matrix without using the autoencoder framework (that is, without applying the random projection and autoencoder steps) with different ensemble sizes. We found that the improvement in clustering performance is diminished in most cases (Figure 3, red boxes), suggesting that the improved clustering performance is due to the random projection and autoencoder steps implemented in the proposed framework in addition to the ensemble step. Notably, the autoencoder framework also enhance the data signal-to-noise ratio in most cases as can been seem from the improved performance of autoencoder-based *k*-means cluster compared to direct *k*-means clustering on the raw input at the ensemble size of 1.

Another interesting observation is that the variance of the clustering outputs in general decrease with the increasing number of base clustering (Figure 3). While the ensemble of *k*-means clustering without random projection and autoencoder does not improve cell type identification accuracy, it does reduce the clustering variability, and therefore, result in more stable and reproducible clustering outputs compared to a single run of *k*-means clustering. These results are consistent with previous findings [21]. In comparison, the ensemble of autoencoder-based clustering lead to both more accurate cell type identification and a reduction of variability, both of which are desirable characteristics for scRNA-seq data analysis.

### Autoencoder-based SIMLR ensemble

While the autoencoder-based cluster ensemble framework is able to improve the performance of a standard *k*-means clustering algorithm in both accuracy and reproducibility of cell type clustering, we wondered if such an ensemble framework could also improve the performance of the latest clustering algorithm. To this end, we applied the proposed framework to a state-of-the-art kernel-based clustering algorithm, SIMLR, designed specifically for cell type identification on scRNA-seq data. Because the computational complexity of SIMLR grows exponentially with the number of cells in a dataset, we focused our evaluation on the two smaller datasets (i.e. GSE84371 and GSE82187). Similar to *k*-means clustering, we found in these cases that the performance of the autoencoder-based SIMLR ensemble improved with increased ensemble size (Figure 5). Clustering variability also generally decreased with larger ensemble sizes. These results demonstrate that the proposed autoencoder-based cluster ensemble framework also leads to more accurate cell type identification and clustering reproducibility from scRNA-seq data for SIMLR.

**Figure 4.**
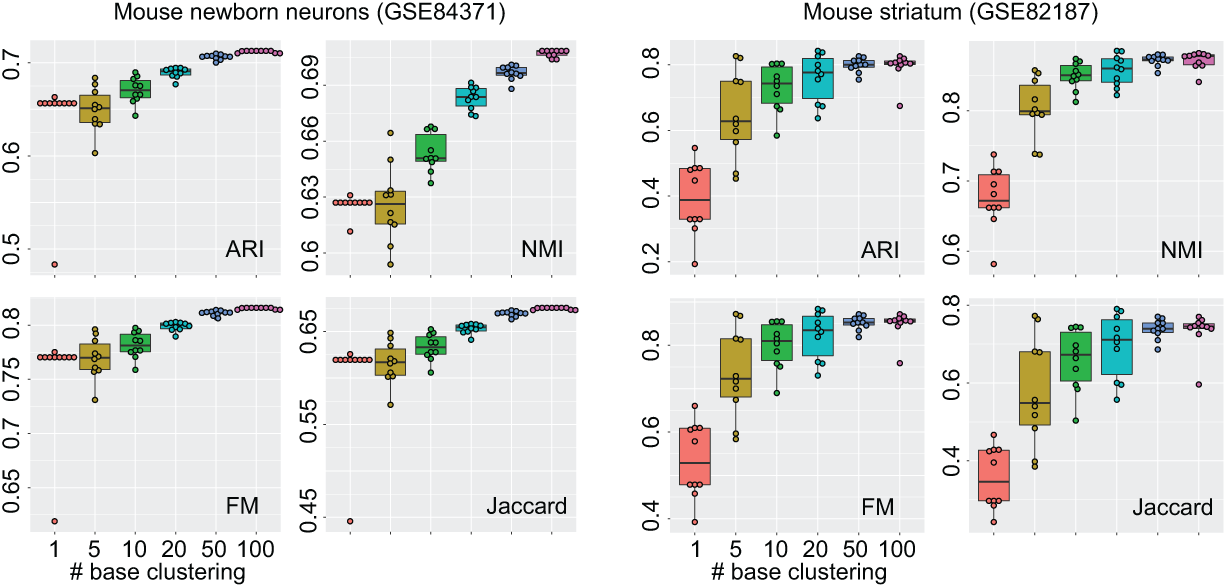
Evaluation of autoencoder-based SIMLR ensemble. Ensemble sizes range from 1 to 100 were tested using four evaluation metrics in two scRNA-seq datasets.

**Figure 5.**
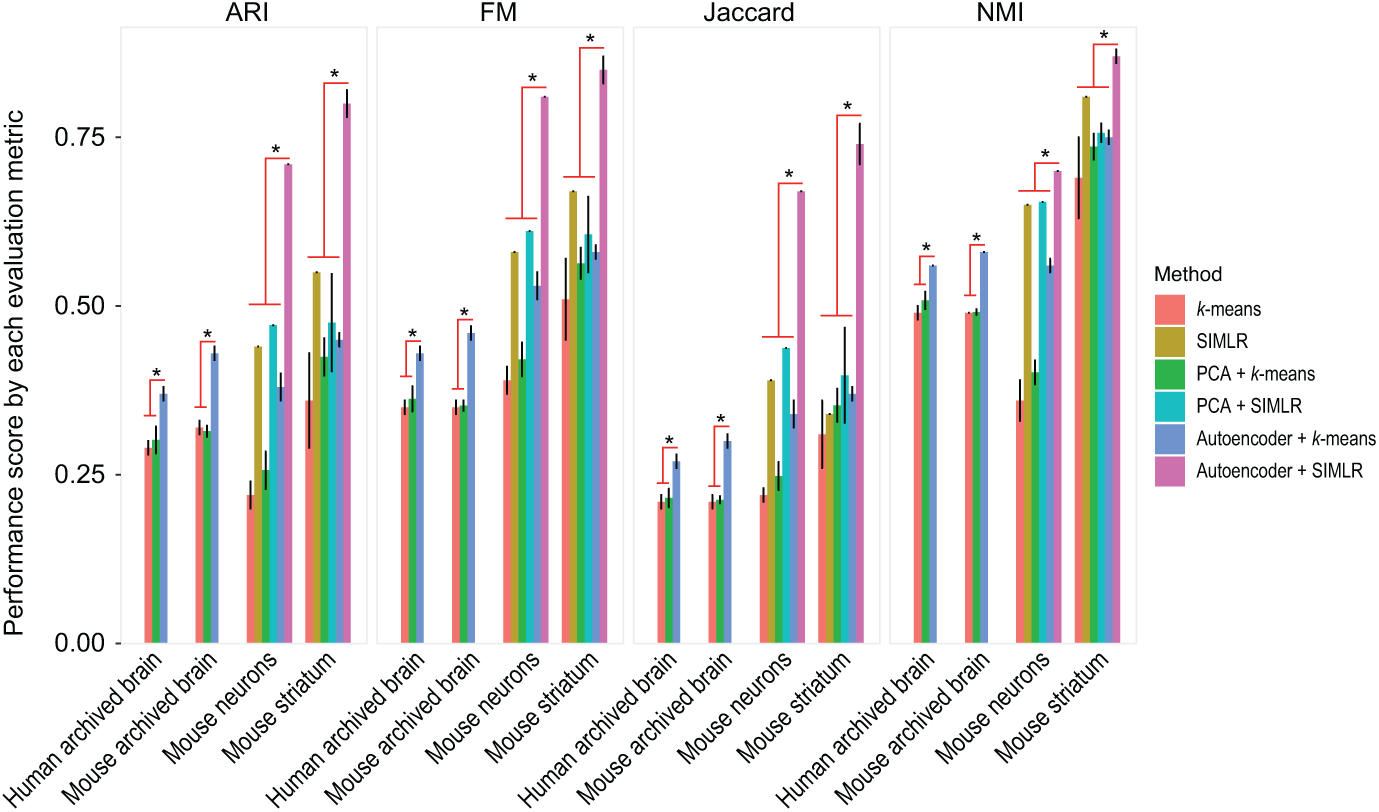
Comparison of autoencoder-based clustering framework with PCA-based dimension reduction and clustering using the four evaluation metrics. Statistical significance (*p <*0.001; denoted by *\**) of either autoencoder with *k*-means clustering against the rest for human and mouse archived brain datasets, or autoencoder with SIMLR clustering against the rest for mouse neurons and striatum datasets were performed using Wilcoxon Rank Sum test (two-sided).

While cell type clustering accuracy improves with the larger ensemble sizes for both *k*-means clustering algorithm and SIMLR (Figure 3 and 5), we observed that this improvement plateaus at an ensemble size of 50 (Figure 5). We therefore recommend an ensemble size of 50 as a good trade-off between clustering output quality and computational time. Note that the computational complexity of the proposed cluster ensemble framework increases linearly with respect to the ensemble size.

### Performance comparison of autoencoder-based cluster ensemble

Typically, a single run of a clustering algorithm is used to identify cell types from a given scRNA-seq dataset. An interesting question is how much improvement the proposed autoencoder-based cluster ensemble offers compared to the common clustering procedure where a clustering algorithm is directly applied to raw gene expression data (that is, without random projection and autoencoder steps). To address this, we next quantified cell type clustering accuracy from the direct application of *k*-means and SIMLR clustering to the raw gene expression input and compared these with the autoencoder-based cluster ensemble of *k*-means and SIMLR, respectively. Note that an ensemble size of 50 was used for the cluster ensemble. *k*-means clustering and the random projection step in ensemble clustering are non-deterministic; while SIMLR contains technically non-deterministic steps (including a *k*-means step), we found that repeated runs on the same raw dataset with different random seeds produced identical clustering partitions. Consequently, SIMLR may be thought of as functionally deterministic. To account for variability in the clustering results for stochastic methods, we repeated clustering ten times and calculated the standard deviation across multiple runs. Table 1 summaries these results.

Specifically, the autoencoder-based *k*-means ensemble improved cell type clustering for an average of about 30% in the four evaluation datasets according to all four evaluation metrics (Table 1). Clustering variability was also typically smaller using the autoencoder-based *k*-means ensemble. Perhaps more strikingly, the cell type clustering accuracy as measured by ARI and Jaccard metrics for autoencoder-based SIMLR ensemble improved about 50% to 100% compared to using SIMLR alone on the raw expression matrix for the mouse neurons and striatum datasets. Moreover, we found that the cell type clustering accuracy of SIMLR is substantially better than the standard *k*-means clustering algorithm, suggesting SIMLR is indeed an effective clustering algorithm for scRNA-seq data analysis. Therefore, the further gain in clustering accuracy by coupling SIMLR with the proposed autoencoder-based cluster ensemble is of practical importance and will add to the state-of-the-art methods for scRNA-seq data analysis.

### Comparison of autoencoder-based cluster ensemble with PCA-based clustering

We next compared the performance of autoencoder-based cluster ensemble with PCA-based clustering. PCA is a commonly used dimension reduction method and has been widely used for reducing the high-dimensionality of scRNA-seq data prior to clustering cell types. By benchmarking the performance of cell type clustering across the evaluation datasets, we found that in almost all cases autoencoder-based clustering ensemble outperformed PCA-based approach for both *k*-means clustering and SIMLR according to all four evaluation metrics (Figure 5). We confirmed the statistical significance of these performance improvements using the Wilcoxon Rank Sum test. These results further demonstrate the utility of the autoencoder-based cluster ensemble for more accurate clustering of cell types in scRNA-seq datasets.

## Discussions

There may be further opportunities to build on the proposed method:

Firstly, we performed hyperparameter optimization over four datasets searching for the most robust configuration. While our chosen configuration was the most consistently accurate over all possible combinations of these four datasets, we saw that it was not among the most accurate configurations for two of the optimization datasets individually. Additionally, there is no guarantee that this combination falls into a global or near-global optimum across scRNA-seq datasets in general, or that such an optimum exists. Devising a way to produce parameter configurations based on individual dataset characteristics without ground-truth labels may be an avenue for further exploration.

Secondly, the current iteration of our proposed method uses random subspace projection to reduce the dimension of datasets prior to autoencoder training. An additional direction for future research may include exploring other methods of basic dimension reduction, such as weighted gene selection based on variability or other metrics; this may be more useful in capturing cell type-specific characteristics by retaining genes containing more biological signal related to cell type.

Lastly, while the clustering algorithms *k*-means and SIMLR were used as independent components in our current proposed framework, an interesting direction for future work might be the development of an artificial neural network architecture and training method which performs simultaneous dimension reduction and clustering. An integrated approach such as this may facilitate the exploration of clustering output in the reduced feature space.

## Conclusions

High throughput scRNA-seq technology is transforming biological and medical research by allowing the global transcriptome profiles of individual cells from heterogeneous samples and tissues to be quantified with high precision. Cell type identification has become essential in scRNA-seq data analysis, and clustering has been the key computational approach used for this task. In this study, we have proposed an autoencoder-based ensemble clustering approach by incorporating several state-of-the-art techniques in a computational framework.

We evaluated the proposed clustering framework on its impact on the level and robustness of cell type identification accuracy using a collection of scRNA-seq datasets with pre-defined cell type annotations. Based on previously defined gold standards for each scRNA-seq dataset, we demonstrate that the proposed framework is highly effective for cell type identification. The application of the proposed framework to both a standard *k*-means clustering algorithm and a state-of-the-art kernel-based clustering algorithms, SIMLR, illustrates its generalisability and applicability to other clustering algorithms. We therefore envision the proposed framework being flexibly adopted into the common workflow for scRNA-seq data analysis.

## Methods

The autoencoder-based cluster ensemble framework is summarised in Figure 1A. The proposed framework accepts scRNA-seq data in the form of an *N* × *M* expression matrix (denoted as *X*) where *N* is the number of cells and *M* is the number of genes.

### Dimension reduction by autoencoders

Genes are randomly selected from the input dataset to produce a set of “random projection” datasets *X*^*t*^ (*t* = 1, *…, T*), each with a dimension of *N* × *M* ′. The purpose of this step is to create ‘diversity’ [27] in subsequent encodings and individual clusterings of these datasets to achieve a more robust consensus in the resulting ensemble. Following the random projection step, each matrix *X*^*t*^ is then used to train a fully-connected autoencoder neural network. An autoencoder is an artificial neural network consisting of two sub-networks: an encoder and a decoder, intersecting at a ‘bottleneck’ layer of a smaller size than the original input. The network is trained to reconstruct the original input with minimum error, forcing the network to learn to encode the information contained within the smaller latent space of the output of the bottleneck layer[30].

In the autoencoders used with our framework, the encoder accepts samples of cell data from *X*′ as input. The encoder contains a single hidden layer and an output layer which produces reduced-dimension encodings of the aforementioned samples. The decoder subnetwork accepts these encoded samples as input, passing these through a single hidden layer and an output layer which produces reconstructions of the original samples. In both subnetworks, the activation function of the hidden layer is a ‘Leaky ReLu’ [31]; linear activation is applied to all other layers.

Each autoencoder is trained by minimising reconstruction error using the mean squared loss function:

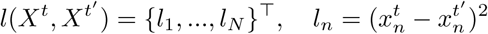

where *X*^*t*^ is the input expression matrix from the *t*^th^ random projection and *X*^*t*′^ is the autoencoder’s reconstruction of *X*^*t*^.

Following training, each matrix *X*^*t*^ is fed through its respective autoencoder and a low-dimension encoded dataset is extracted from the encoder output. Training and hyperparameter optimisation of autoencoders are discussed in Section 4.1.

### Clustering algorithms

To perform clustering on dimension-reduced datasets generated from autoencoders, we utilised both a standard *k*-means clustering algorithm with Lloyd’s implementation [39] and a kernel-based clustering algorithm (SIMLR) specifically designed for scRNA-seq data analysis [10].

Given an initial set of random centres *m*_1_, *…, m*_*K*_, and a distance matrix *D* (typically computed from Euclidean space), the algorithm first finds the closest cluster centres for each of all cells based on their expression profiles 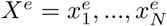:

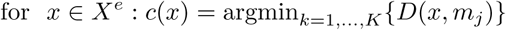

and then updates the cluster centres:

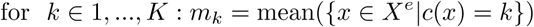

The output is the assignment of each cell based on its expression profile *x* to a cluster *k ∈* 1, *…, K*.

SIMLR calculates the distance matrix for cells using multiple kernels as follows:

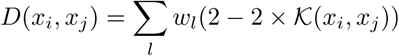

where *w*_*l*_ is the weight of a Gaussian kernel function for a pair of cells defined as follows:

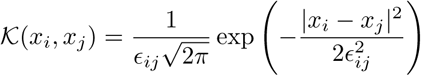

where *ϵ*_*ij*_ is the variance and *|x*_*i*_ *− x*_*j*_*|*^2^ is the squared distance between cell *i* and *j*, calculated from their expression profiles *x*_*i*_ and *x*_*j*_. To test the proposed framework, we utilised SIMLR (Version 1.8.0) implemented in Bioconductor (Release 3.7).

The number of clusters to be created was set according to the number of predefined cell types/classes in each scRNA-seq data for both the *k*-means clustering and SIMLR (see Section 10). After obtaining individual clustering outputs (denoted as *P* ^*t*^) from either *k*-means clustering or SIMLR, a fixed-point algorithm for obtaining hard least squares Euclidean consensus partitions was applied to compute the consensus *P*_*E*_ of individual partitions [28]:

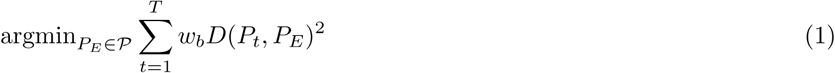

in which *w*_*b*_ is the weight associated with individual clustering output and is set to 1 in our case, and *D*(*P*_*t*_, *P*_*E*_)^2^ is the squared Euclidean function for computing the distance of an individual partition with the consensus partition.

Together, the proposed autoencoder-based cluster ensemble framework can be summarised in pseudocode as below.

### Data Description and Evaluation

This section summarises the scRNA-seq datasets and performance assessment metrics utilised for method evaluation.

#### Algorithm 1 Autoencoder-based cluster ensemble

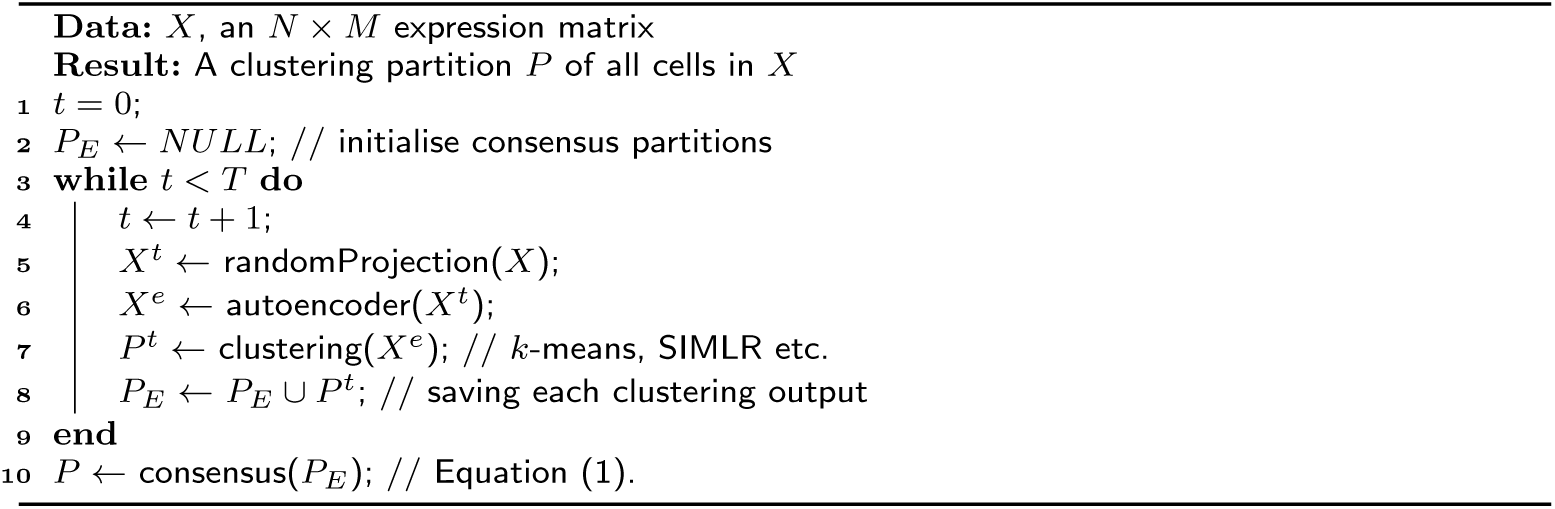

#### scRNA-seq datasets

A collection of eight publicly available scRNA-seq datasets (Table 2) were utilized in this study. These datasets were downloaded from the NCBI GEO repository, the EMBL-EBI ArrayExpress repository, or the Broad Institute Single-Cell database portal. The *log*_2_-transformed transcripts per million (TPM) or counts per million (CPM) values (as determined by the original publication for a given dataset) were used to quantify full length gene expression for datasets generated by SMARTer or SMART-seq2 protocols. UMI-filtered counts were used to quantify gene expression for the Drop-seq datasets. All datasets have undergone cell-type identification using biological knowledge from their respective original publications which we retain for evaluation purposes. For each dataset, genes detected in less than 20% of cells were removed. This step trims the number of genes and allows only those that are expressed in at least a subset of cells to be considered for subsequent analyses. Four datasets were used to optimise autoencoder hyperparameters. We present evaluation benchmarking results for four additional datasets.

#### Evaluation metrics

A common approach to assess the performance of clustering methods for cell type identification in scRNA-seq data analysis is to compare the concordance of the clustering outputs of cells with a ‘gold standard’. As mentioned above, such a gold standard may be obtained from orthogonal information such as cell type marker genes and/or other biological knowledge of cell type characteristics. In this study, cell type annotations from their original publications are used as ‘gold standards’. For each dataset, number of clusters for both *k*-means clustering and SIMLR were set as the number of pre-defined classes based on its original publication and the concordance between the clustering outputs and the ‘gold standard’ were measured using different metrics. Here we employed a panel of four evaluation metrics including Adjusted Rand index (ARI), normalized mutual information (NMI), Fowlkes-Mallows index (FM), and Jaccard index [40] (Figure 6).

**Figure 6.**
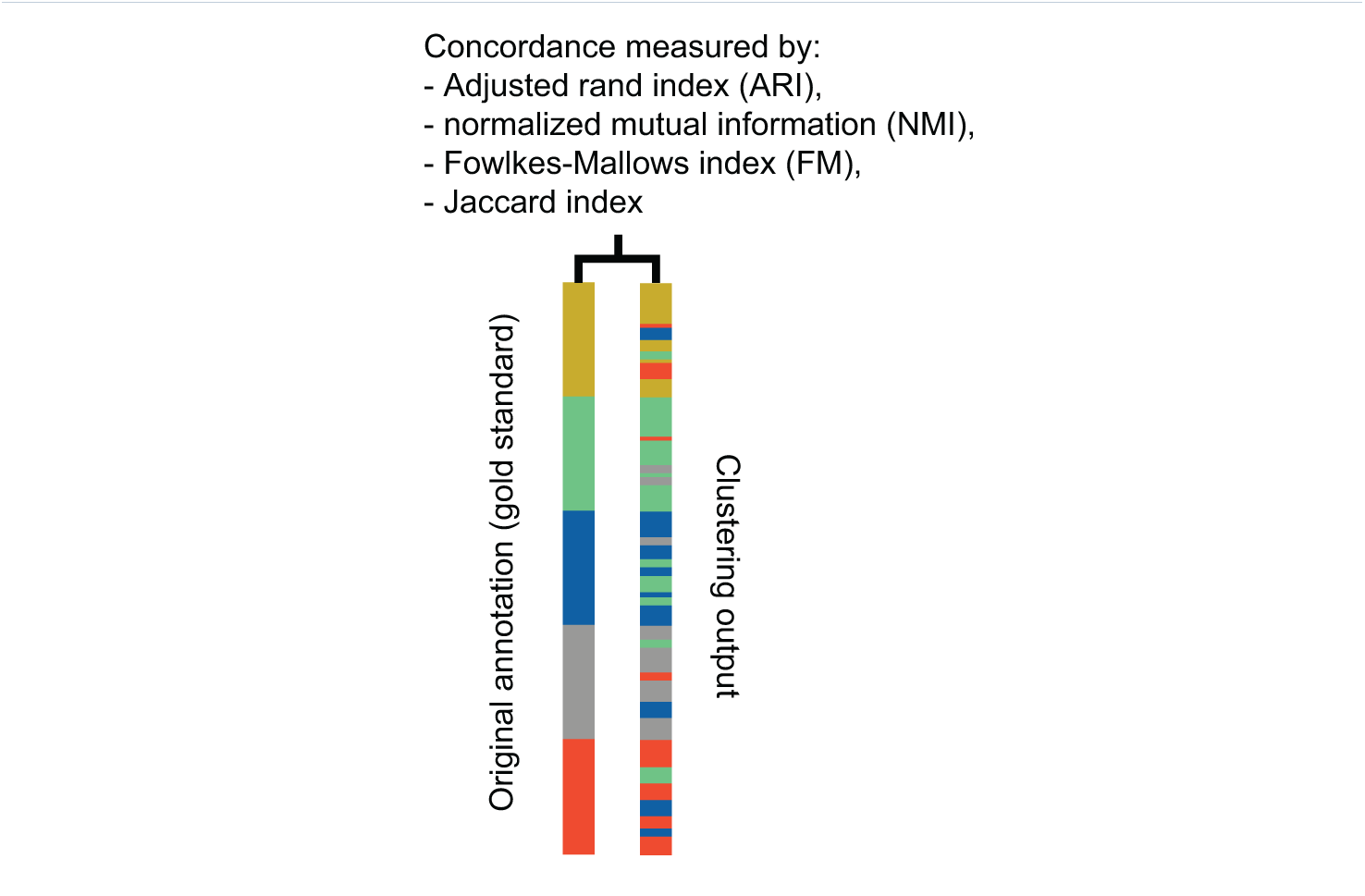
A schematic showing the quantification of concordance of the clustering output with the original ‘gold standard’ annotation using a panel of evaluation metrics.

Let *G, P* be the cell partitions based on the gold standard and the clustering output respectively. We define *a*, the number of pairs of cells assigned to the same group in both partitions; *b*, the number of pairs of cells assigned to the same cell type in the first partition but to different cell types in the second partition; *c*, the number of pairs of cells assigned to different cell types in the first partition but to the same cell type in the second partition; and *d*, the number of pairs of cells assigned to from different cell types in both partitions. ARI, FM, and Jaccard index can then be calculated as follows:

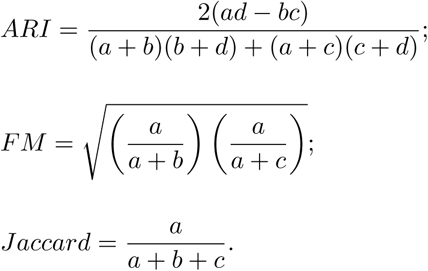

Let *G* = *{u*_1_, *u*_2_, *…, u*_*k*_*}* and *P* = *{v*_1_, *v*_2_, *…, v*_*k*_*}* denote the gold standard and the clustering partition across *K* classes, respectively. NMI is defined as follows:

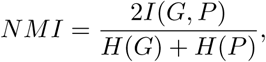

where *I*(*G, P*) is the mutual information of *G* and *P*, defined as

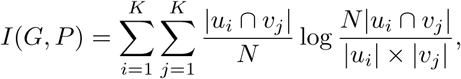

and *H*(*G*) and *H*(*P*) are the entropy of partitions *G* and *P* calculated as

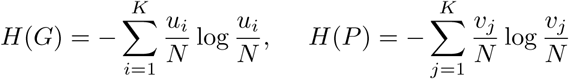

where *N* is the total number of cells.

## List of abbreviations

scRNA-seq: Single-cell RNA-seq;
PCA: principal component analysis;
(tSNE): t-distributed stochastic neighbour embedding;
ARI: adjusted rand index;
FM: Fowlkes-Mallows index;
(SIMLR): kernel-based clustering algorithm;
(CPM): counts per million;
(TPM): transcripts per million.

## Declarations

### Ethics approval and consent to participate

Not applicable.

### Consent for publication

Not applicable.

### Availability of data and material

The datasets generated and/or analysed during the current study are available from the GEO, EBI, and Broad repositories. Details of datasets are listed in Table 1.

### Authors’ contributions

P.Y. and T.A.G. conceived the study. T.A.G. led the experimental design and data analyses. T.K. contributed to the data curation and analyses; L.N. contributed to the algorithm design with input from D.T.; P.Y. and T.A.G. interpreted the experimental results with input from J.G.B. and J.Y.H.Y.; P.Y. and T.A.G. wrote the manuscript with input from all authors. All authors reviewed and approved the final version of the manuscript.

## Acknowledgements

The authors thank their colleagues at the School of Mathematics and Statistics; and School of Life and Environmental Sciences for informative discussion and valuable feedback.

